# *GABRA1* frameshift variants impair GABA_A_ receptor proteostasis

**DOI:** 10.1101/2024.11.28.625971

**Authors:** Marnie P. Williams, Ya-Juan Wang, Jing-Qiong Kang, John H. Perryman, Ting-Wei Mu

## Abstract

The gamma-aminobutyric acid type A receptor (GABA_A_R) is the most common inhibitory neurotransmitter-gated ion channel in the central nervous system. Pathogenic variants in genes encoding GABA_A_R subunits can cause receptor dysfunction and lead to genetic epilepsy. Frameshift variants in these genes can result in a premature termination codon, producing truncated receptor subunit variants. However, the pathogenic molecular mechanism as well as functional implications of these frameshift variants remains inadequately characterized. This study focused on four clinical frameshift variants of the α_1_ subunit of GABA_A_R (encoded by the *GABRA1* gene): K401fs (c.1200del), S326fs (c.975del), V290fs (c.869_888del), and F272fs (c.813del). These variants result in the loss of one to three transmembrane helices, whereas wild type α1 has four transmembrane helices. Therefore, these variants serve as valuable models to evaluate membrane protein biogenesis and proteostasis deficiencies of GABA_A_Rs. In HEK293T cells, all four frameshift variants exhibit significantly reduced trafficking to the cell surface, resulting in essentially non-functional ion channels. However, the severity of proteostasis deficiency varied among these four frameshift variants, presumably due to their specific transmembrane domain deletions. The variant α_1_subunits exhibited endoplasmic reticulum (ER) retention and activated the unfolded protein response (UPR) to varying extents. Our findings revealed that these frameshift variants of *GABRA1* utilize overlapping yet distinct molecular mechanisms to impair proteostasis, providing insights into the pathogenesis of GABA_A_R-associated epilepsy.

## Introduction

Gamma-aminobutyric acid type A receptors (GABA_A_Rs) are ligand-gated ion channels that bind to GABA, the primary inhibitory neurotransmitter in the brain. Thus, GABA_A_Rs are a major contributor to inhibitory signaling in the central nervous system. This inhibitory signaling is critical in maintaining the excitation-inhibition (E-I) balance, which is essential for normal brain activity (1). Therefore, dysfunction of GABA_A_Rs consequently results in impaired inhibitory signaling, causing an E-I imbalance which leads to seizures and epilepsy disorders (2). Pathogenic variants of genes encoding GABA_A_R subunits which disrupt proper channel function are among the prominent causes of genetic epilepsy.

For proper function, the GABA_A_R subunits must fold into their native structure and assemble as a heteropentameric protein in the endoplasmic reticulum (ER) (3). To date, 19 different subunits are known in mammals, including α_1-6_, β_1-3_, γ_1-3_, θ, ε, π, δ, and ρ1-3 (4, 5). The most common GABA_A_R subtype found in the mature mammalian brain is comprised of two α_1_ subunits, two β_2_ subunits, and one γ_2_ subunit (6). Once assembled, GABA_A_Rs are trafficked to the surface membrane, where the GABA neurotransmitter binds at the interface between the α and β subunits (7). Upon GABA binding, fast inhibitory chloride currents hyperpolarize the postsynaptic neuron to inhibit neuronal firing (5).

Variant subunits may present as misfolded proteins in the ER, preventing efficient GABA_A_R trafficking and function. Terminally misfolded proteins must be cleared promptly, through proteasomal or lysosomal degradation (8–10). Additionally, the presence of misfolded GABA_A_R subunits in the ER may activate the unfolded protein response (UPR), which attempts to restore protein homeostasis (11). The UPR regulates ER proteostasis through three pathways, namely, the ATF6 (activating transcription factor 6), IRE1 (inositol-requiring enzyme 1), and PERK (protein kinase R-like ER kinase) pathways (11–13). Upon activation, ATF6 travels from the ER to the Golgi, and its N-terminal cytosolic fragment (ATF6-N) is cleaved and translocates to the nucleus to act as a transcription factor and enhance the expression of downstream genes, such as BiP and ERdj3 (14). IRE1 activation results in the splicing of XBP1 (XBP1s), while PERK activation phosphorylates the eukaryotic translation initiation factor 2’s α-subunit (eIF211) (15, 16). Although PERK is the primary inducer of CHOP, a pro-apoptotic transcription factor, IRE1 activation can also lead to the activation of CHOP (15, 17). Gaining insight into the cellular mechanisms involved in adapting to misfolded proteins is crucial to uncovering the impact of the variant subunits on GABA_A_R function.

To date, over 1,000 variants, such as missense, frameshift, and nonsense, observed in genes encoding GABA_A_R subunits, have been reported in the NIH ClinVar (ncbi.nlm.nih.gov/clinvar). While the pathogenicity of missense variants has been predicted (18), and a fraction of them has been characterized (2), research on frameshift and nonsense variants remains limited. Here, we focused on four frameshift variants in the *GABRA1* gene: K401fs425X (c.1200del), S326fs328X (c.975del), V290fs299X (c.869_888del), and F272fs287X (c.813del), which are regarded as K401_fs_, S326_fs_, V290_fs_ and F272_fs_, respectively, in this study. These frameshift variants, caused by single nucleotide deletions, introduce premature termination codons (PTCs), leading to truncated variant α_1_ subunits. In this study, we characterized the trafficking, degradation, UPR activation, and function of these four *GABRA1* frameshift variants.

## Results

### Molecular features and phenotypes of four GABRA1 frameshift variants

The four *GABRA1* frameshift variants result in truncated α_1_ subunits to various degrees due to the introduction of PTCs. Protein sequence alignment with the wild type (WT) α_1_ subunit demonstrated the residues adjacent to the termination site for each frameshift variant **(Fig. 1A)**. F272 is located at the end of transmembrane helix 1 (TM1), V290 is in the middle of TM2, S326 is near the middle of TM3, whereas K401 is situated in the long cytosolic loop between TM3 and TM4 (**Fig. 1A**, **1B**). Membrane topology prediction suggested that K401_fs_ and S326_fs_ retained three TM helices, V290_fs_ has one or two TM helices, and F272_fs_ has only one TM helix, compared to the four TM helices in the WT α_1_ subunit (**Fig. 1C**)(19). These frameshift variants likely cause substantial structural defects on the α_1_ subunit, leading to receptor dysfunction and consequently epilepsy.

**Figure 1.**
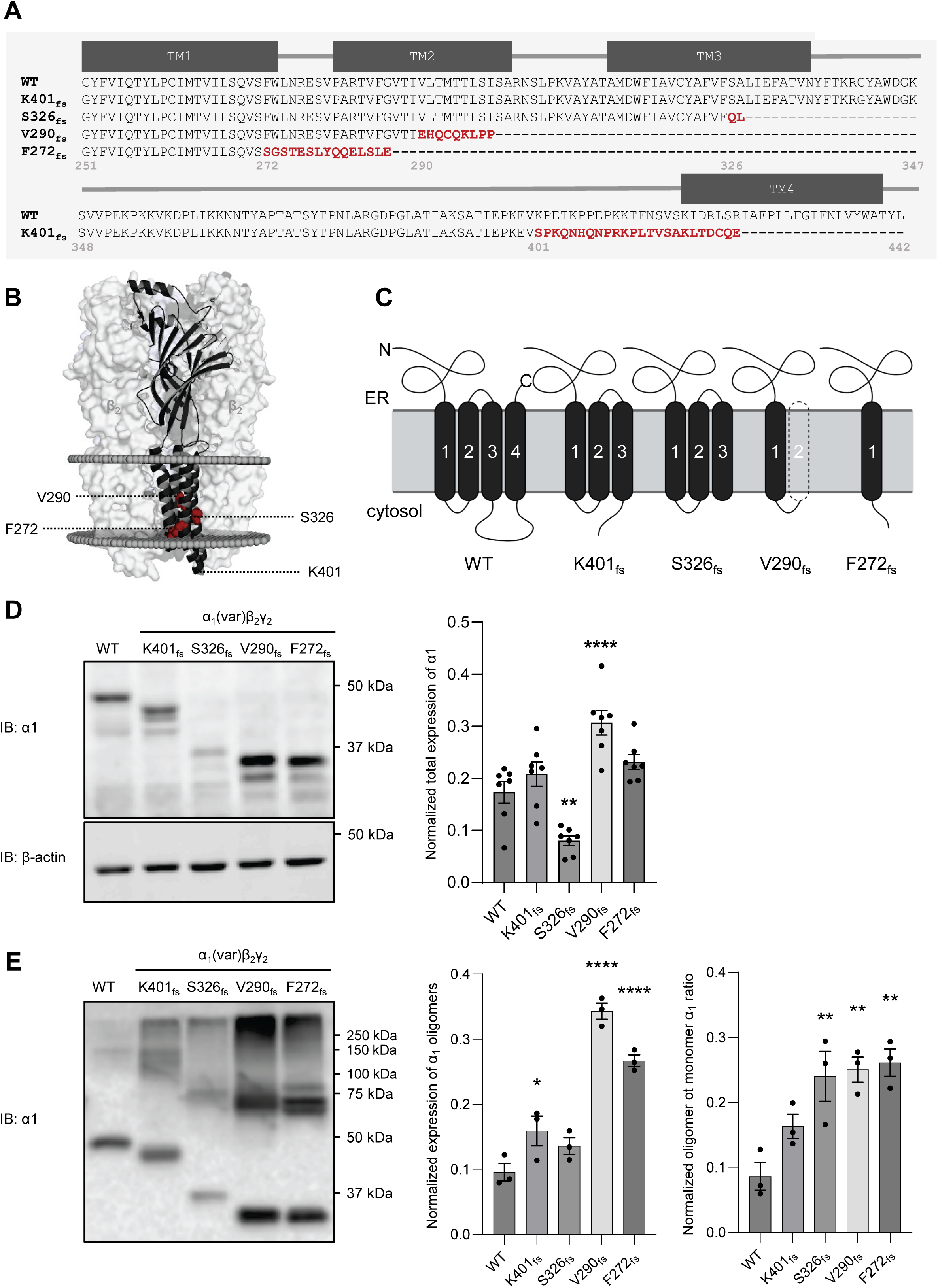
*GABRA1* frameshift variants result in transmembrane domain deletions and distinct total expression levels of the α_1_ subunit. (**A**) Primary sequence alignments, focusing on the transmembrane domains, were shown of the WT and the *GABRA1* frameshift variants: K401fs425X (c.1200del), S326fs328X (c.975del), V290fs299X (c.869_888del), and F272fs287X (c.813del), which are regarded as K401_fs_, S326_fs_, V290_fs_ and F272_fs_, respectively, in this study. (**B**) Cartoon representation of the positions of the four frameshift variants in the cryo-EM structure of the α_1_ subunit, built from 6D6T.pdb using PyMOL. (**C**) Schematic representation of the wild-type (WT, left) α_1_ subunit membrane topology and the predicted transmembrane domain deletions of the α_1_variants resulted from the *GABRA1* frameshift variants. (**D**) The α_1_ subunits (WT or variant) were co-transfected with β_2_ and γ_2_ subunits at a 1:1:1 ratio in HEK293T cells. Cells were harvested and total proteins were extracted 48 hours after transfection, followed by SDS-PAGE and Western blot to detect the α_1_ subunit. The bands of the WT α_1_ subunit and K401_fs_, S326_fs_, V290_fs_ and F272_fs_ variants, were detected at 48, 42, 35, 32, 30 kDa, respectively. Protein band intensity was quantified with ImageJ and shown on the right (n=7). β-actin was used as a loading control. (**E**) Transfected HEK293T cells with α_1_ (WT or variant), β_2_, and γ_2_ subunits were harvested and lysed 48 hours post-transfection. A non-reducing SDS-PAGE gel and western blotting was used to analyze the oligomerization of the variant α_1_ subunits compared to WT. Bands detected above the 50 kDa marker corresponded to the oligomerized forms of α_1,_ while bands detected below the 50 kDa marker corresponded to the monomeric α_1_ subunits. Quantification of total oligomer expression and oligomeric / monomeric α1 was shown on the right (n =3). * *p*< 0.05; **, *p* < 0.01; ****, *p* < 0.0001.

Clinical features of these four *GABRA1* frameshift variants were summarized in **Table 1**. A patient presenting with infantile-onset refractory epileptic encephalopathy was found to have a *de novo* frameshift variant of *GABRA1*, K401_fs_ (**Table 1**) (20). This patient also exhibited severe developmental delay; however, it remains unclear whether delay stems from the *GABRA1* variant or the patient’s coexisting condition, Williams-Beuren syndrome (WBS) (20). In another case, a patient diagnosed with childhood absence epilepsy (CAE) harbored the *de novo GABRA1* frameshift variant, S326_fs_, yet responded well to treatment with valproate, an anti-seizure drug (**Table 1**) (21). Additionally, the NIH Clinvar reported that another patient with CAE carried the pathogenic *GABRA1* F272_fs_ variant, but no further details were provided.

**Table 1.** Clinical features of *GABRA1* frameshift variants of K401_fs_, S326_fs_, V290_fs_, and F272_fs_ in the GABA_A_R.

| **Variant** | **Condition** | **Classification by ClinVar** | **Reference** |
| --- | --- | --- | --- |
| K401<sub>fs</sub> | Infantile-onset refractory epileptic encephalopathy | Pathogenic | (20) |
| S326<sub>fs</sub> | Childhood absence epilepsy (CAE) | Pathogenic | (21) |
| V290<sub>fs</sub> | Developmental and epileptic encephalopathy | Pathogenic/Likely pathogenic | Scarlett's GABRA1 Village |
| F272<sub>fs</sub> | Childhood absence epilepsy (CAE) | Pathogenic | NIH ClinVar |

Furthermore, a patient diagnosed with global development delay was found to carry the *de novo GABRA1* frameshift variant, V290_fs_ (**Table 1**). This patient met developmental milestones through the 6-month checkup; however, at 9 months old, the patient exhibited severe motor delays. At 12 months old, the patient was referred for Early Intervention evaluation and services due to delays in gross and fine motor skills. By 15 months old, speech delay was noted, and the patient was referred to neurology. Dysmorphic features were also observed, leading to a referral to genetics. A brain MRI (magnetic resonance imaging) revealed asymmetry of the lateral ventricles but was otherwise normal. The dysmorphic features noted by the geneticist included a prominent forehead, narrow midface, syndactyly, small hands and feet, and lateral displacement of the inner canthi. At 2 years old, the patient demonstrated brief episodes of eye rolling and twitching, which were initially sporadic. However, routine and video electroencephalograms (EEGs) were normal, therefore, anticonvulsants or anti-epileptic drugs were never prescribed. Despite the high risk for seizures associated with this variant, the patient has remained seizure-free thus far.

### GABRA1 frameshift variants display varying total protein levels, decreased surface trafficking, and reduced function of GABA_A_Rs

Since proteostasis deficiency is a major disease-causing mechanism for GABA_A_R variants (22, 23), we evaluated the effect of the four frameshift variants on GABA_A_R protein expression, surface trafficking, and function. To determine protein expression levels of WT versus variant α_1_ subunits: K401_fs_, S326_fs_, V290_fs_, and F272_fs_, the α_1_ subunits were co-transfected with β_2_ and γ_2_ subunits at a 1:1:1 ratio in HEK293T cells. Cells were harvested 48 hours after transfection and lysed, and total proteins were subjected to SDS-PAGE and western blot analysis to visualize total α_1_ protein expression. We observed varying total α_1_ protein levels: compared to WT, K401_fs_ did not significantly affect α_1_protein levels, S326_fs_ led to a significant 2.2-fold protein level reduction, V290_fs_ resulted in a significant 1.7-fold protein level increase, and F272_fs_ caused a 1.3-fold protein level increase although this increase was not statistically significant (**Fig. 1D**). Consistent with our observation, S326_fs_ has been previously reported to display a reduced protein expression compared to WT (24). Interestingly, the V290_fs_ variant subunit led to significantly increased total α1 expression, possibly due to the introduction of an extra cysteine (C293) (**Fig 1A**) which could potentially form more stable complexes such as 11_1_dimers through crosslinking compared to WT. Furthermore, under non-reducing conditions, we observed substantially increased α1 dimers for V290_fs_ (at 64 kDa) and F272_fs_ (at 60 kDa) and their higher-order oligomers compared to WT (**Fig 1E**). Moreover, the ratio of the oligomeric to monomeric α1 was increased in the S326_fs_, V290_fs_, and F272_fs_ variants compared to WT (**Fig 1E**). These results suggest that the V290_fs_ and F272_fs_ variants’ excessive oligomerization compared to WT could lead to the detrimental accumulation of misfolded proteins within the cell.

Surface expression is critical for GABA_A_R function since the ion channel must reach the plasma membrane to conduct currents. Therefore, we determined the surface expression of these α_1_ variants using surface biotinylation and immunocytochemistry (**Fig. 2A**, **2B**). 48 hours post-transfection of WT or variant α_1_ with β_2_ and γ_2_ subunits in HEK293T cells, a biotinylation assay was performed to extract surface proteins (**Fig. 2A**). Both surface and cytosolic proteins were processed by SDS-PAGE and Western blot to detect α_1_ expression, with Na^+^/K^+^ ATPase and β-actin serving as loading controls for surface and cytosolic proteins, respectively. In all the surface fractions (even lanes in **Fig. 2A**), no β-actin was detected, indicating that indeed during the surface biotinylation assay, no cytosolic contaminants were introduced to the surface protein pools. The normalized surface to cytosolic α_1_expression was dramatically decreased for all four variants compared to WT: 22% for K401_fs_, 29% for S326_fs_, 27% for V290_fs_, and 13% for F272_fs_ (**Fig. 2A**). Additionally, immunocytochemistry was used to visualize surface and intracellular expression of the α_1_ subunits. Surface α_1_ subunits were stained with an anti-α_1_ antibody with an epitope recognizing its extracellular region and visualized with Alexa-488 (green) **(Fig. 2B**, third row). Intracellular α_1_ was then stained following membrane permeabilization and visualized with Alexa-568 (red) **(Fig. 2B**, second row). Consistent with the biotinylation results, the normalized surface to cytosolic α_1_ fluorescence intensity ratio was dramatically reduced for all four variants (**Fig. 2B**).

**Figure 2.**
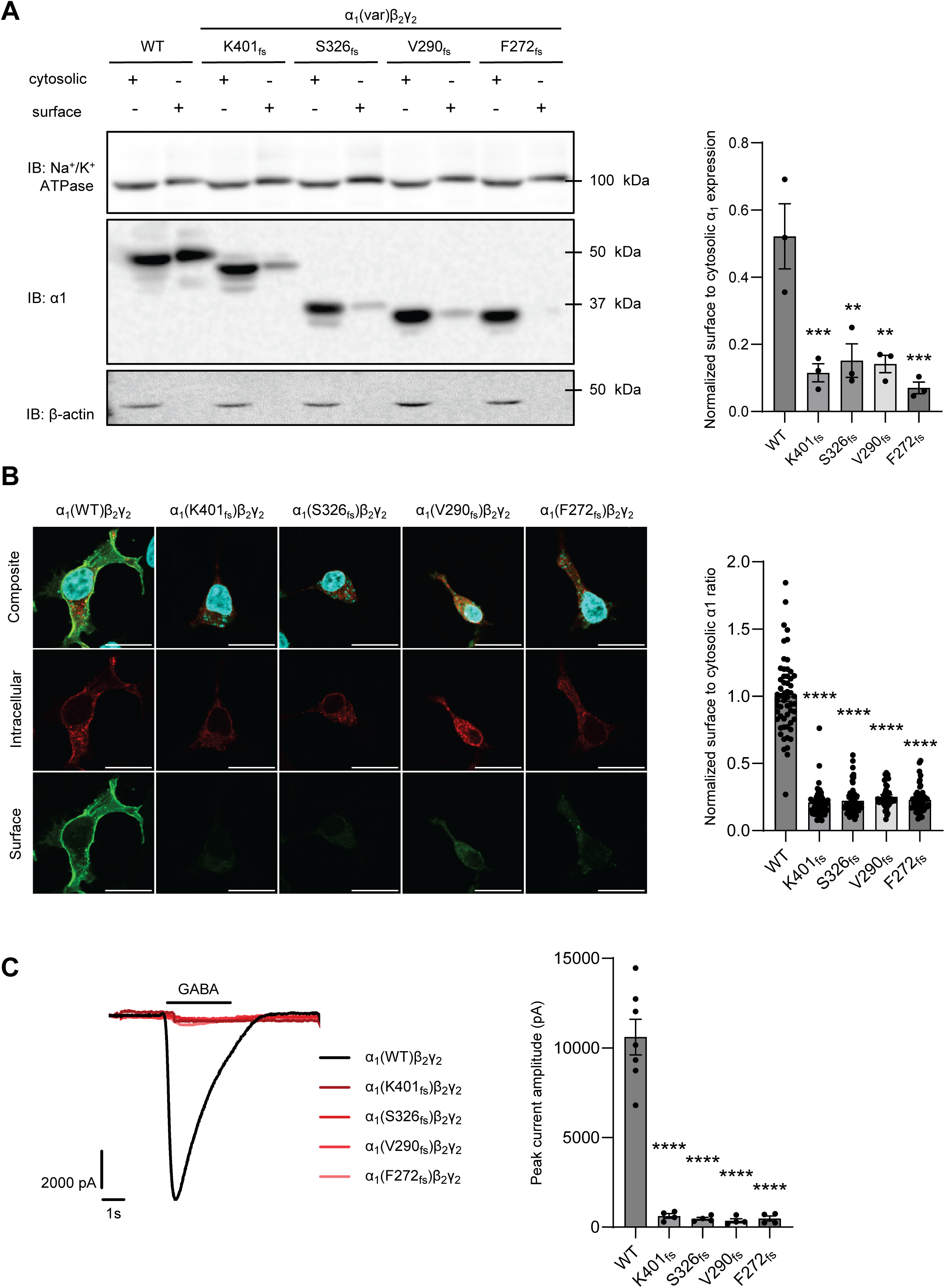
*GABRA1* frameshift variants resulted in decreased surface expression and function of GABA_A_R. HEK293T cells were transfected with α_1_subunits (WT or variant), β_2_, and γ_2_subunits at a 1:1:1 ratio. (**A**) The biotinylation assay was carried out 48 hours after transfection to extract surface proteins. Both surface and cytosolic proteins were processed by SDS-PAGE and Western blot to detect the α_1_ subunit. Protein band intensity was quantified with ImageJ and the normalized surface to cytosolic α1 band ratio was shown on the right (n=3). Na^+^/K^+^ ATPase was used as a loading control for surface proteins, and β-actin as a loading control for cytosolic proteins. (**B**) Cells were fixed with 4% formaldehyde without membrane permeabilization and incubated with an anti-α_1_ antibody. A secondary antibody with Alexa-488 (green) was applied to represent particularly the surface α_1_. Afterwards, we carried out membrane permeabilization, and incubated the cells with the anti-α_1_ antibody and further with a secondary antibody with Alexa-568 (red) to label the intracellular α_1_. The fluorescence intensity was analyzed using ImageJ and the quantification is shown on the right (scale bar = 20 μm, n=40-57). (**C**) Automated patch-clamping was used to record GABA_A_R current at a holding potential of -60 mV, 48 hours after transfection. GABA was applied at 100 μM for 3 s and the peak current amplitudes were acquired and plotted (n=4-7 of ensemble recordings; each ensemble recording enclosed 20 cells). Data is presented as mean ± SEM. ANOVA followed by Dunnett’s test was used for statistical analysis. ** *p* < 0.01; ***, *p* < 0.001; ****, *p* < 0.0001.

We further evaluated the effect of the α1 variants on GABA-induced currents using automated patch-clamping electrophysiology in HEK293T cells transfected with WT or variant α_1_ along with β_2_ and γ_2_ subunits. In response to saturating GABA conditions, the average peak current amplitudes for recombinant GABA_A_Rs containing WT, K401_fs_, S326_fs_, V290_fs_ and F272_fs_ α_1_ subunits was 10607 ± 990.5, 622 ± 128.7, 467.3 ± 78.71, 360.5 ± 110.9, and 482.8 ± 138.7 pA, respectively (**Fig. 2C**). Therefore, the decreased surface expression in all four α_1_variants, which is critical for functional GABA binding, led to over 90% reduction of GABA-induced peak currents compared to WT.

These results demonstrated that all four frameshift variants substantially reduced surface trafficking, leading to the loss of function of GABA_A_Rs. However, interestingly, these variants have differing steady-state protein levels, suggesting their context-dependent impact on protein stability (**Fig. 1D**).

### GABRA1 frameshift variants result in α_1_ subunit ER retention and distinct protein stability

With the decreased surface expression of the α_1_ variant subunits compared to WT, we investigated whether these variants are retained in the ER. First, we performed an endoglycosidase H (endo H) assay to quantify trafficking efficiency from the ER to the Golgi apparatus (**Fig. 3A**). Endo H digestion cleaves glycan groups modified in the ER, while more complex glycan groups attached in the Golgi apparatus are resistant to endo H digestion. Peptide-N-Glycosidase F (PNGase F), which can cleave both glycan groups that are installed in the ER and Golgi, served as a control for non-glycosylated forms of the α_1_ variants (**Fig. 3A**, lanes 3, 6, 9, 12, and 15). The bottom bands after endo H digestion, aligning with PNGase F treated bands, represented endo H-sensitive α1 subunits, whereas the top bands correspond to endo H-resistant α_1_ subunits (**Fig. 3A**, lanes 2, 5, 8, 11, and 14) (25, 26). The ratio of endo H resistant to total α_1_ was used to quantify the trafficking efficiency of α_1_. All α_1_ variants demonstrated a decreased ratio of endo H resistant to total α1 after endo H digestion (**Fig. 3A**, cf. lanes 5, 8, 11, and 14 to lane 2): K401_fs_ showed a reduction to approximately 40%, S326_fs_ to 20%, V290_fs_ to 23%, and F272_fs_ to 22% compared to WT (**Fig. 3A**). These results indicated that all four variants substantially reduced the ER-to-Golgi trafficking efficiency of the α_1_ subunit.

**Figure 3.**
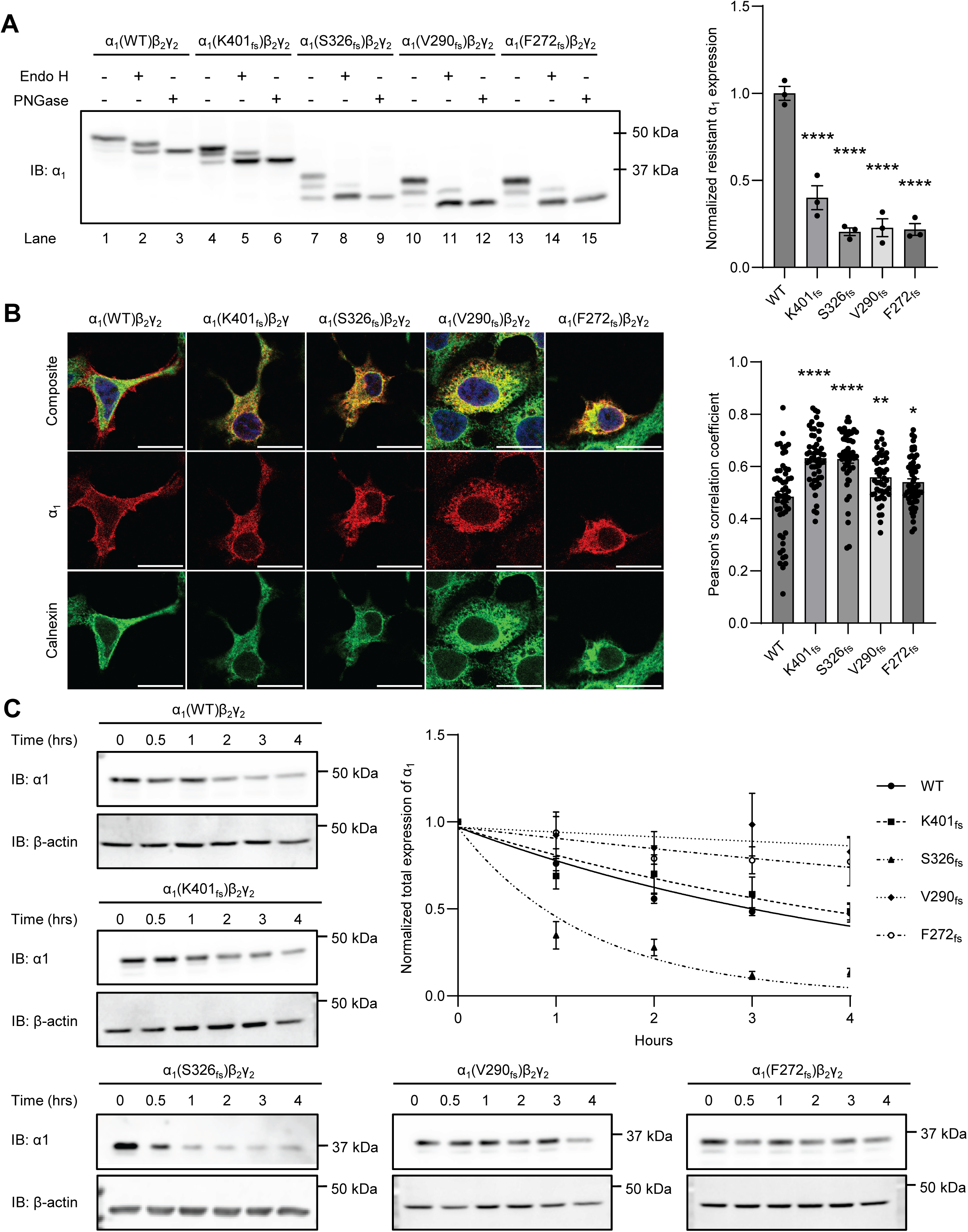
*GABRA1* frameshift variants result in the α_1_ subunit’s retention in the ER and varied protein stability. (**A**) Using an Endo-H assay, the α_1_ variants demonstrated reduced Endo-H resistance compared to WT. PNGase F digestions generate non-glycosylated α1, as in lanes 3, 6, 9, 12, and 15. After Endo-H digestions, top bands are the resistant bands, whereas bottom bands, which align with PNGase F-digested α1 bands, are the sensitive bands, as in lanes 2, 5, 8, 11, and 14. Quantification of the normalized Endo-H resistant α1, as a measure of the ER-to-Golgi trafficking efficiency, is shown on the right (n=3). (**B**) Co-localization was determined using immunocytochemistry staining for both the α_1_ subunit (green) and a known ER-resident protein, Calnexin (red). The Pearson’s coefficient was analyzed using ImageJ and quantification is shown on the right (n= 30-56) (**C**) A cycloheximide-chase assay was performed to determine α_1_’s stability of GABA_A_Rs harboring the frameshift variants of K401_fs_, S326_fs_, V290_fs_ and F272_fs_ (n=3-8). A one-phase decay non-linear regression was used to determine half-life. Data is presented as mean ± SEM. ANOVA followed by Dunnett’s test was used for statistical analysis. *, *p* < 0.05; ****, *p* < 0.0001.

To further confirm ER retention, we analyzed the co-localization of α_1_ variants with calnexin, an ER-resident type I transmembrane protein, using immunofluorescence staining. All four α_1_ frameshift variants displayed substantial overlap with calnexin staining and exhibited greater Pearson’s correlation coefficients compared to WT (**Fig. 3B**). Consistent with our observation, K401_fs_ has been previously reported to display increased ER localization compared to WT (27). The co-localization of the α_1_ variants with an ER protein coupled with the lack of ER-to-Golgi trafficking demonstrate the significant ER retention of the α_1_ variants. This retention contributes to the decreased surface expression and function of the pathogenic GABA_A_Rs.

Misfolding-prone membrane proteins retained in the ER can lead to the formation of aggregates or rapid removal, resulting in differing protein stability and steady-state protein levels. We used a cycloheximide (CHX)-chase assay to assess α_1_ protein stability (**Fig. 3C**). Following transfection, HEK293T cells were treated with cycloheximide, a potent protein synthesis inhibitor to measure the degradation of α_1_ variants. We observed varying α_1_ protein stability among the four α_1_ frameshift variants: compared to WT (t_1/2_ = 3.14 hrs), the K401_fs_ variant exhibited a slightly slower degradation rate (t_1/2_ = 3.83 hrs), the S326_fs_ variant demonstrated much faster α_1_ degradation (t_1/2_ = 0.92 hr), and the V290_fs_ and F272_fs_ variants showed significantly increased stability (t_1/2_ > 4 hrs) (**Fig. 3C**). The distinct protein stability of the four variants correlates with their steady-state protein levels observed earlier, suggesting that more stable variants like V290_fs_ and F272_fs_ accumulate and potentially form aggregates in the ER.

### GABRA1 frameshift variants are subject to proteasomal or lysosomal degradation and nonsense-mediated decay

To evaluate the proteolytic degradation pathway of terminally misfolded α_1_ variants, we treated transfected HEK293T cells with bafilomycin A1 (BafA1) and MG132 to inhibit lysosomal and proteasomal degradation, respectively. Accordingly, treatment with MG132 (200 nM, 24 hrs) led to the accumulation of ubiquitinated proteins (**Fig. 4A, Supplemental Fig. S1A**), indicating effective inhibition of the proteasome, whereas BafA1 treatment (20 nM, 24 hrs) resulted in increased LC3b isoform II and the LC3b II/I ratio (**Fig. 4A**, **Supplemental Fig. S1B**), indicating efficient inhibition of the lysosomal degradation pathway. Here, we find that the S326_fs_ α_1_ variant, which has the fastest degradation rate, displayed increased α_1_ protein expression following treatment with BafA1 or MG132 (**Fig. 4A**), indicating that the S326_fs_ α_1_variant utilized both the proteasome and lysosome for clearance. Interestingly, the F272_fs_ variant was shown to utilize the lysosomal pathway, however, the V290_fs_ variant exhibited only a slight increase in protein expression after lysosomal inhibition (**Fig. 4A**). The V290_fs_ variant’s substantial protein accumulation may mask any changes in expression following degradation inhibition. Additionally, under our experimental conditions, the K401_fs_ variant showed no significant change in α_1_ expression following treatment with either BafA1 or MG132 (**Fig. 4A**).

**Figure 4.**
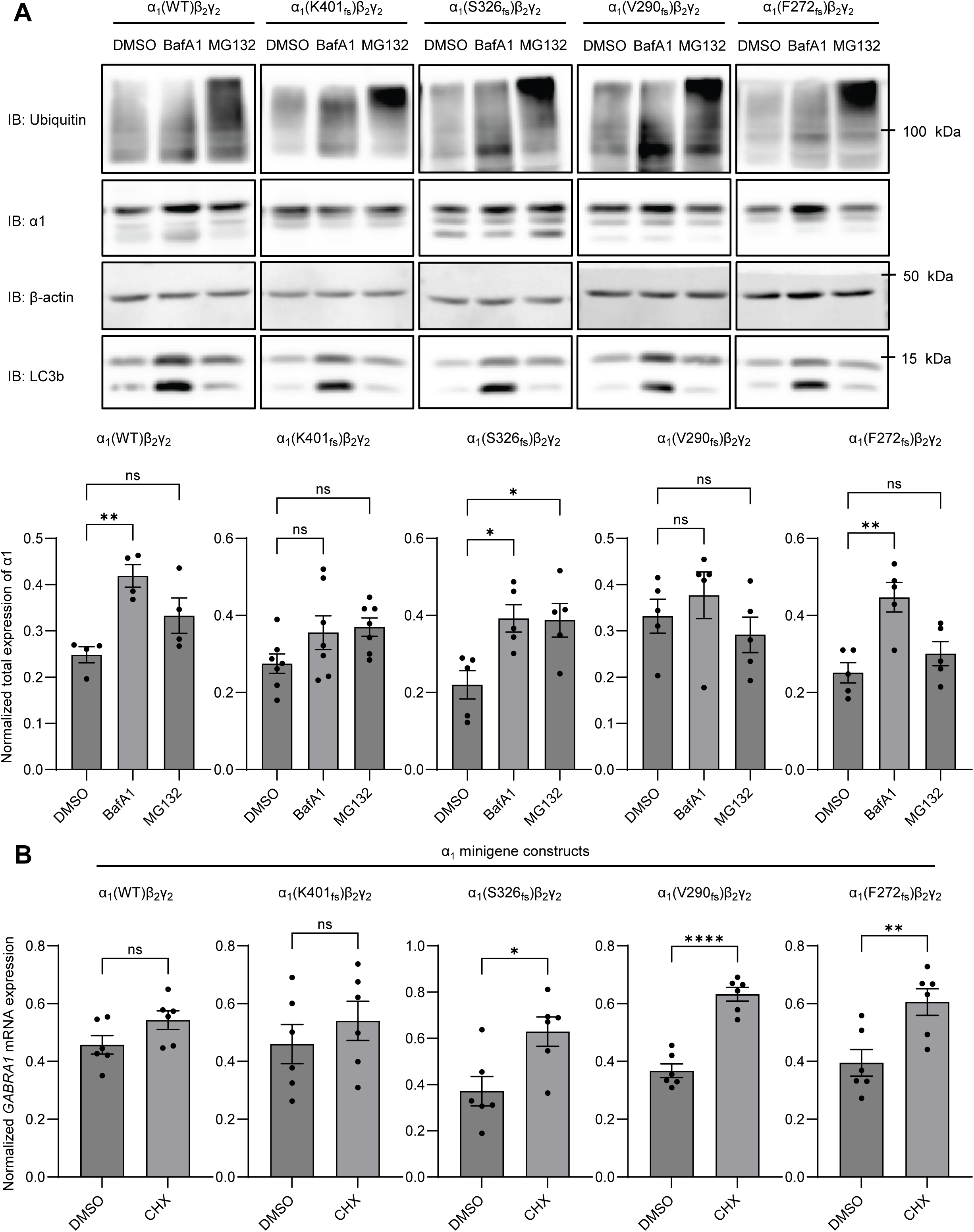
*GABRA1* frameshift variants utilize mRNA and protein degradation mechanisms. (**A**) HEK293T cells were transfected similar to **Figure 1-2**; MG132 (200 nM, 24 hrs) and BafA1 (20 nM, 24 hrs) were used to inhibit proteasomal and lysosomal degradation, respectively. Total α_1_ proteins were extracted 48 hours post transfection, followed by SDS-PAGE and Western blot to detect the α_1_ protein. Protein band intensity was quantified with ImageJ and shown on the bottom (n=4). β-actin was used as a loading control. (**B**) WT and variant α_1_ subunit minigenes were constructed by introducing intron 8 (1242 bp) into the cDNA in order to study NMD. The α_1_ minigenes were co-transfected with β_2_ and γ_2_ subunits at a 1:1:1 ratio in HEK293T cells. Cycloheximide (CHX) was added as an inhibitor of NMD at 100 µg/ml for 6 hrs, 48 hours after transfection. Using qRT-PCR, relative mRNA *GABRA1* expression was quantified (n=6). Data is presented as mean ± SEM. Unpaired t-test was used for comparison of two groups, and ANOVA followed by Dunnett’s test was used for comparison of more than two groups. *, *p* < 0.05; **, *p* < 0.01; ***, *p* < 0.001; ****, *p* < 0.0001.

In addition to proteostasis defects, we investigated whether the four frameshift variants were subjected to nonsense-mediated decay (NMD), an mRNA degradation mechanism. During translation, NMD arrests mRNAs that contain premature termination codons (PTCs) in proximity and upstream to an exon-exon junction, and then degradation machinery is recruited and results in mRNA decay (28). We transfected HEK293T cells with α_1_ minigene constructs derived from human *GABRA1* cDNAs containing the entire intron 8 (**Supplemental Fig. S1C**), enabling the activation of NMD mechanisms (24). Splicing of this construct will produce an exon-exon junction between exon 8 and 9, which is necessary for the recruitment of the NMD machinery (24). To determine NMD susceptibility, HEK293T cells were treated with cycloheximide (CHX), a known indirect NMD inhibitor that inhibits protein translation and consequently, inhibits NMD following transfection (24, 29). After treatment with CHX (100 μg/ml, 6 hrs) we find that only the K401_fs_ variation is not subject to NMD, likely due to the PTC occurring much later in the protein compared to the other variants thereby, making it less recognizable to NMD machinery (**Fig. 4B**). Although the S326_fs_, V290_fs_, and F272_fs_ variants are subjected to NMD, approximately 20% of mRNA transcripts are still able to escape NMD and undergo protein translation (24, 30).

### The GABRA1 frameshift variants activate the UPR differently

Since we demonstrated that the four frameshift variants result in ER retention, we aimed to determine whether these variants induce UPR activation, a major ER stress response mechanism. The UPR responds to stress by activating three cellular signaling pathways, namely, ATF6, IRE1, and PERK. We first evaluated the effect of the four variants on the ATF6 activation. The mRNA expression of hallmark ATF6 downstream targets, including BiP (gene: *HSPA5*) and ERdj3 (gene: *DNAJB11*), was quantified using real-time quantitative reverse transcription polymerase chain reaction (qRT-PCR). Thapsigargin (Tg), a potent UPR activator, was administered to HEK293T cells as a positive control for UPR activation (**Supplemental Fig. S2A**). Compared to WT, aggregation-prone V290_fs_ and F272_fs_, increased the mRNA levels of BiP and ERdj3 significantly, whereas K401_fs_ and S326_fs_ did not have significant influence (**Fig. 5A**), indicating that these variants have distinct effects on the ATF6 pathway. Furthermore, we determined that the mRNA expression of the pro-apoptotic ER stress marker, CHOP, was upregulated significantly for K401_fs_ and V290_fs_ compared to WT (**Fig. 5A).**

**Figure 5.**
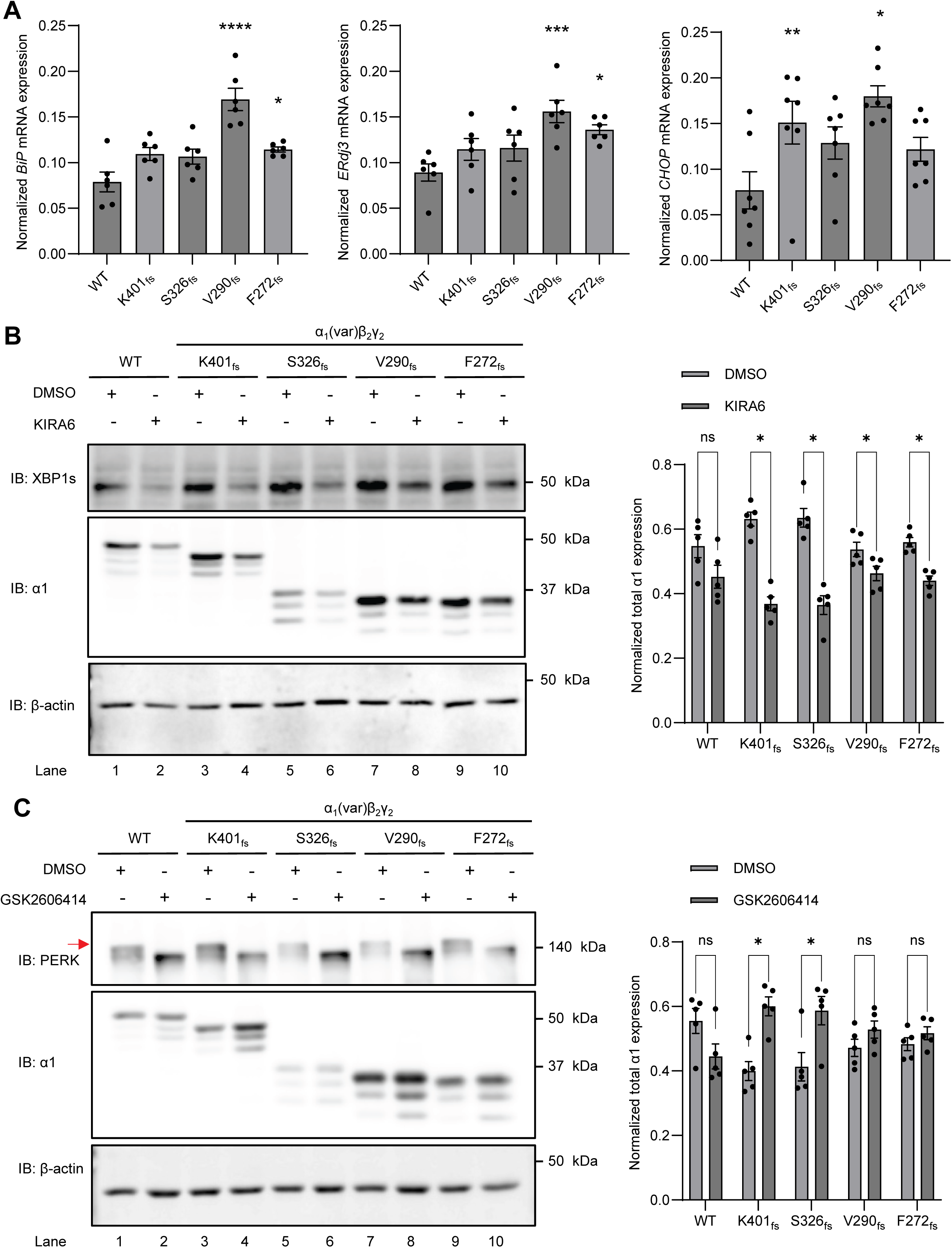
*GABRA1* frameshift variants activate the UPR to variable extents. (**A**) Relative mRNA expression of UPR targets were analyzed, including BiP and ERdj3 (downstream targets of the ATF6 arm), as well as CHOP, a pro-apoptotic transcription factor (n=6-7). (**B**) A potent IRE1/XBP1-s inhibitor, KIRA6 (1 µM, 24 hrs), was applied to HEK293T cells expressing WT or frameshift α_1_ variants. Post 24 hrs of treatment, cells were harvested and total proteins were extracted, followed by SDS-PAGE and Western blot to detect the proteins of interest. Protein band intensity was quantified with ImageJ and shown on the right (n=5). (**C**) A potent PERK inhibitor, GSK2606414 (0.5 µM, 24 hrs), was applied to HEK293T cells expressing WT or frameshift α_1_ variants. Post 24 hrs of treatment, cells were harvested and total proteins were extracted, followed by SDS-PAGE and Western blot to detect the proteins of interest. The activated phosphorylated PERK was labeled by the red arrow, which was used for quantification. Protein band intensity was quantified with ImageJ and shown on the right (n=5). β-actin was used as a loading control. Data is presented as mean ± SEM. ANOVA followed by Dunnett’s post-hoc test was used for statistical analysis. *, *p* < 0.05; **, *p* < 0.01; ***, *p* < 0.001; ****, *p* < 0.0001.

In addition to the ATF6 pathway activation following ER stress, IRE1 activation results in its dimerization and autophosphorylation. This activates its RNase domain in the cytosol, leading to the splicing of XBP1. The spliced form of XBP1 (XBP1s) is translocated into the nucleus to induce the expression of molecular chaperones to enhance the ER folding capacity. Western blot analysis revealed that compared to WT, the S326_fs_, V290_fs_, and F272_fs_ variants displayed a slight increase in untreated XBP1s levels (**Fig. 5B**, cf. lanes 3, 5, 7, and 9 to lane 1), suggesting these variants mildly enhanced the IRE1/XBP1s pathway. Inhibition of IRE1 using KIRA6 (1μM, 24h) reduced the XBP1s protein levels substantially and resulted in a unanimous reduction of the detergent-soluble fractions of the four α_1_ variants (**Fig. 5B**, **Supplemental Fig. S2B**). These results highlight a critical role of the IRE1/XBP1s pathway in maintaining proteostasis of these α_1_ variants. Moreover, we evaluated the effect of the four variants on the induction of the PERK pathway by analyzing PERK phosphorylation since PERK dimerizes and auotophosphorylates in response to ER stress. Western blot analysis demonstrated that compared to WT, all four variants did not change untreated PERK phosphorylation significantly, indicating that these variants did not promote the PERK pathway (**Fig. 5C**, cf. lanes 3, 5, 7, and 9 to lane 1). We then applied a PERK pathway inhibitor, GSK2606414 (0.5 µM, 24 hrs), to HEK293T cells and found that inhibiting PERK slightly increased α_1_ protein levels of the K401_fs_ and S326_fs_ variants, but not the V290_fs_, and F272_fs_ variants (**Fig. 5C**).

These results revealed that the V290_fs_ and F272_fs_ α_1_ variants, which likely form large aggregates and exhibit more stability than WT, activate ATF6 and IRE1 pathways to manage their proteostasis deficiencies. Additionally, the K410_fs_ and V290_fs_ variants induce greater pro-apoptotic stress than WT through induction of CHOP. Furthermore, the variants displayed distinct changes in protein expression upon PERK inhibition, likely due to their distinct topology-based protein folding defects. However, findings from IRE1 inhibition suggest a shared ER stress response mechanism among the four frameshift variants.

## Discussion

**Figure 6** illustrates our proposed characterization of the frameshift variants in the α_1_ subunit (*GABRA1* gene) of the GABA_A_R. The variants S326_fs_, V290 _fs_, and F272 _fs_, but not K401_fs_, are subjected to NMD. Once translated, all the four α_1_ variants misfold and are retained in the ER, leading to their inability to traffic to the Golgi and further to the surface membrane and loss of function. Thus, these variants induce ER stress and activate the UPR to various degrees. Their ER retention also leads to impaired degradation of the variant subunit. Interestingly, the V290_fs_ and F272_fs_ variants are more stable than WT, likely forming large aggregates in the ER, and targeted to the lysosome for clearance; both variants active IRE1/XBP1s arm and ATF6 arm of the UPR to handle misfolded proteins. Compared to WT, the destabilized S326_fs_ variant leads to much faster degradation kinetics and utilizes both the lysosome and proteasome for its efficient removal.

**Figure 6.**
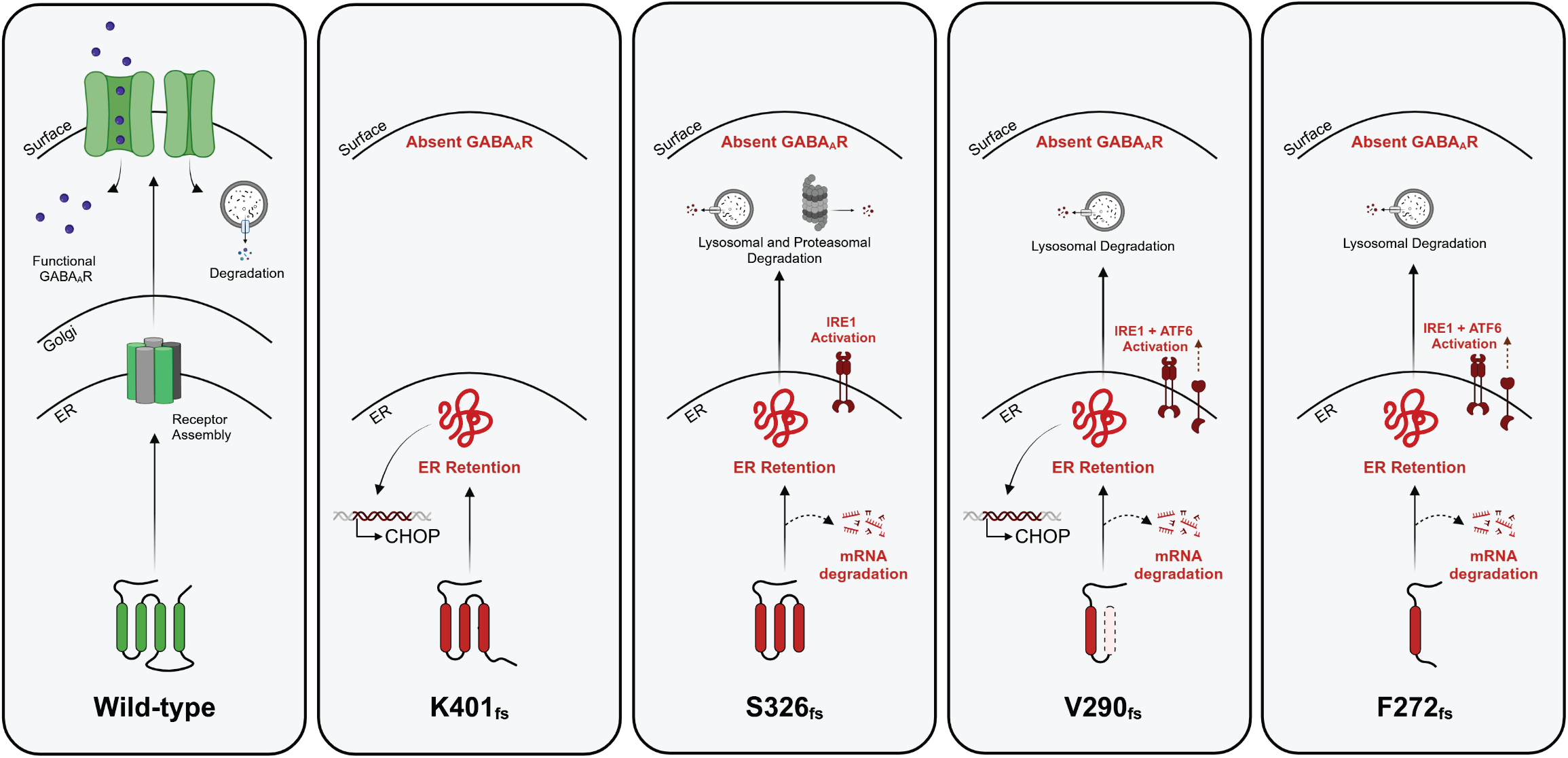
Proposed mechanism for the *GABRA1* frameshift variants in the GABA_A_R. The α_1_ frameshift variants K401_fs_, S326_fs_, V290_fs_ and F272_fs_ lead to ER retention and inefficient trafficking to cell plasma membrane, and thus decreased function. Furthermore, due to their distinct transmembrane domain deletions, folding capability and stability, they are subject to nonsense mediated decay (NMD), proteasomal and/or lysosomal degradation and unfolded protein response (UPR) activation, specifically the ATF6 and IRE1/XBP1s pathways.

Our data determined that the K401_fs_, S326_fs_, V290_fs_ and F272_fs_ variants result in trafficking deficiencies and therefore, have decreased ionotropic receptor function. Due to their distinct transmembrane domain deletions, these variants exhibit variable expression levels, which are influenced by their oligomerization, specific proteolytic degradation capability, protein stability, and UPR activation (**Fig. 1-5**). However, these variants unanimously show reduced ER-to-Golgi trafficking and thus, decreased α_1_ surface expression compared to WT (**Fig. 2 and 3**). As a result, the variants resulted in over a 90% reduction of GABA-induced peak current using whole-cell patch clamping (**Fig. 2C**), which is consistent with their known pathogenicity (**Table 1**). Furthermore, we observed a significant increase in co-localization of α_1_ and calnexin (**Fig 3B**) compared to WT, indicating defective anterograde trafficking and degradation as the subunits remain retained in the ER.

The misfolded subunits trapped in the ER resulted in protein stability correlating to their α_1_ expression level. Both WT and K401_fs_ exhibited similar α_1_ expression and stability, while the S326_fs_ displayed lower expression and instability (**Fig. 1D**, **3C**). In contrast, the V290_fs_ and F272_fs_ variants resulted in accumulated and stable α_1_ expression (**Fig. 1D**, **3C**). Our data suggest that the cell addresses these variants through mRNA and proteolytic degradation. Specifically, the variants S326_fs_, V290 _fs_, and F272 _fs_ are subjected to nonsense-mediated decay, while the K401_fs_ variant escaped this mRNA surveillance mechanism (**Fig. 4B**). Furthermore, the S326_fs_, V290_fs_, and F272_fs_ variants are degraded via the lysosomal pathway, with S326_fs_ also involving the proteasomal pathway (**Fig. 4A**). The ability of the S326_fs_ variant to utilize both degradation pathways likely facilitates the efficient clearance of the misfolded subunit. Interestingly, although K401_fs_ bypasses NMD, it did not show a significant preference for either the lysosomal or proteasomal pathways (**Fig. 4A**, **4B**). These findings warrant further investigation into the protein handling mechanisms associated with these variants.

Excessive ER retention coupled with differential protein processing of these variants by the cell can disrupt proteostasis and lead to ER stress. In response, the cell activates the UPR to restore ER homeostasis and relieve ER stress (11, 31). Our results demonstrate that the V290_fs_ and F272_fs_ variants led to increased BiP mRNA expression, a key chaperone that is critically involved in regulating ER homeostasis and activated the ATF6 arm of the UPR pathway (32). In contrast, although the α_1_variant S326_fs_ was found to be ER retained, it did not result in upregulation of ERdj3, BiP or CHOP (**Fig. 5A**). This result could be due to rapid clearance of the S326_fs_ variant by both the proteasome and lysosome (**Fig. 4A**). Interestingly, both K401_fs_ and V290_fs_ variants exhibited increased levels of CHOP, despite the lack of BiP upregulation in the K401_fs_ variant (**Fig. 5A**). It is also important to note that V290_fs_ and F272_fs_ are predicted to lose at least two transmembrane helices and the intracellular loop between TM3 and TM4, which plays a role in protein interactions with cytosolic factors as well as post-translational modifications, such as phosphorylation and ubiquitination (**Fig. 1C)** (33). These structural defects may compromise beneficial protein-protein interactions and thus, require assistance from the ATF6 pathway to direct their proper protein folding or clearance.

The frameshift variants in the α_1_ subunit of GABA_A_Rs led to varying extents of UPR activation likely due to their differing expression levels, stability, and degradation mechanisms. To further explore the role of the UPR in regulating the expression of the variant α_1_ subunits, we used a known PERK pathway inhibitor, GSK2606414, to block the phosphorylation of eIF2α, a critical mediator of the downstream signaling of PERK (34). Our results showed that PERK inhibition led to a slight increase in the expression of the variants, most notably in the K401_fs_ and S326_fs_ variants (**Fig. 5C**). PERK activation typically inhibits protein synthesis to relieve the protein load in the ER and alleviate ER stress (35). By contrast, blocking this pathway resulted in higher expression of the misfolded subunits, likely by counteracting the stress response mechanism. This suggests that the PERK pathway may play a role in limiting the accumulation of these variants.

We additionally investigated the role of the IRE1 pathway using a known inhibitor, KIRA6, which blocks the RNase activity of IRE1 (15, 31). Our results demonstrate that IRE1 inhibition led to a reduction in detergent-soluble α_1_ fractions for all the variants (**Fig. 5B**), which could be attributed to several effects of IRE1 inhibition such as decreased folding capability in the ER and reduced downstream targets, such as RAMP4 and PDI-P5, both of which are involved in membrane protein stabilization and maturation (36). Moreover, compromised protein folding machinery induced by IRE1 inhibition could result in more misfolded proteins, triggering intervention and compensation by other UPR arms or cellular processes to clear these proteins (37, 38). The role of each UPR arm is much more intricate than simply inhibiting protein synthesis as there is ample crosstalk among the UPR signaling pathways and other protein homeostasis mechanisms. Our results suggest that the frameshift variants each result in a distinct regulation of α_1_ compared to WT, due to their distinct membrane topology and protein sequence.

Given the complex role of the UPR in protein misfolding and accumulation, it is also important to consider how the ER adapts to handle the increased load of retained proteins. It has been reported that the ER has the potential to expand its membrane structure and volume to handle ER-retained proteins, as demonstrated using high-resolution electron microscopy (EM) (39–41). In a study where HeLa cells were exposed to the secretory heavy chain of immunoglobulin M (µ_s_), an ER stressor, the ER volume was found to have increased 2-3 fold during 3 days of µ_s_ induction. Electron micrographs exhibited darker, electron-dense areas, which was achieved by introducing the APEX (pea peroxidase) to the ER using an N-terminal signal peptide and a C-terminal KDEL (39–41). A similar EM study of ER volume changes upon expression of the α_1_ variants merits further investigation.

The WT α_1_ subunit along with the frameshift variants together, which range from one to four TM helices (**Fig. 1C**), serve as valuable transmembrane client proteins to study the molecular mechanisms of membrane protein insertion into the lipid bilayer, particularly with the assistance of molecular chaperones. Recently the EMC complex, especially EMC3 and EMC6, and an ER luminal chaperone Hsp47 have been demonstrated to facilitate the folding and assembly of the α_1_ subunit of GABA_A_Rs (42, 43). Furthermore, it appears that the α_1_ TM4 helix is inserted post-translationally by the EMC (44). Therefore, the stepwise truncation of the four TM helices afforded by these α_1_ frameshift variants could be used to investigate the co-translational or post-translational insertion processes for clinically relevant multipass transmembrane proteins.

There is growing evidence suggesting that an emerging disease-causing mechanism of epilepsy involves protein misfolding, which disrupts ionotropic receptor trafficking and function (31, 45, 46). Moreover, many GABA_A_R variants are resistant to current anti-seizure drugs, which primarily target surface-expressed receptors and fail to address underlying trafficking deficiencies (47). However, proteostasis regulators such as stress-independent UPR activators (45), FDA-approved drugs like dinoprost and dihydroergocristine (48), and GABA_A_R-specific pharmacological chaperones including hispidulin and TP003 (23, 49) have been demonstrated to restore the surface trafficking and function of pathogenic GABA_A_R variants. Thus, further research into enhancing proteostasis as a strategy to rescue these pathogenic receptor variants holds promise for the development of novel therapeutic approaches for epilepsy.

## Experimental procedures

### Chemicals

GSK3606414 (#HY-18072, MedChemExpress) was used for PERK inhibition at 0.5 μM for 24 hours. KIRA6 (#HY-19708, MedChemExpress) was used for IRE1 inhibition at 1 μM for 24 hours. Thapsigargin (Tg; #BML-PE180-0001, Enzo Life Sciences) was used as a positive control for ER stress and UPR activation at 1 μM for 6 hours. MG132 (#A2585, ApexBio) was used for proteasome inhibition at 200 nM for 24 hours. Bafilomycin A1 (BafA1; #11038, Cayman Chemical) was used for lysosome inhibition at 20 nM for 24 hours. Cycloheximide (CHX; #ALX-380-269, Enzo Life Sciences) was used as an indirect NMD inhibitor at 100 μg/mL for 6 hours.

### Plasmids

The pCMV6 plasmids were purchased from Origene, including human GABA_A_R α1 subunit (Uniprot #: P14867-1) (# RC205390), β2 subunit (isoform 2, Uniprot #: P47870-1) (#RC216424), γ2 subunit (isoform 2, Uniprot #: P18507-2) (#RC209260), and pCMV6 Entry Vector plasmid (pCMV6-EV) (#PS100001). The minigene construct of the *GABRA1* cDNA with the inclusion of intron 8 was generated previously (24). The frameshift variants of K401fs425X (c.1200del), S326fs328X (c.975del), V290fs299X (c.869_888del), and F272fs287X (c.813del) in the GABA_A_R α1 subunit were constructed using a QuikChange II site-directed mutagenesis Kit (Agilent Genomics, #200523), and the cDNA sequences were confirmed by DNA sequencing.

### Cell culture and transfection

HEK293T cells (Abgent #CL1032) were cultured and passaged in Dulbecco’s Modified Eagles Medium (DMEM/High Glucose; #SH3024301, Fisher Scientific) with 10% fetal bovine serum (FBS; #SH3039603HI, Fisher Scientific) and 1% penicillin-streptomycin (#SV30010, Fisher Scientific). Cells from passages 15 to 35 were utilized and seeded at 250,000 cells per 6-well and transfected 24 hours after plating with plasmids using *Trans*IT-2020 (#MIR 5404, Mirus Bio) and Opti-MEM^TM^ (#31985070, ThermoFisher). Subunits α_1_β_2_γ_2_ were transfected at a 1:1:1 ratio (i.e. 0.25 μg of plasmid per subunit in a 6-well plate). Then, cell lysates were harvested 48 hours after transfection.

### Western blotting

Cells were harvested by cell scraping, centrifuged at 5000 x g for 2 min, and washed twice with 1x DPBS (#SH3002803, Fisher Scientific). HEK293T cells were lysed using lysis buffer made with 1x TBS (50 mM Tris, pH 7.5, 150 mM NaCl), 2 mM *n*-Dodecyl-β-D-maltoside (DDM) (#DDM5, GoldBio), supplemented with complete protease inhibitor cocktail (#4693159001, Roche), then cells were vortexed and sonicated for 40 seconds each, repeated thrice. PhosSTOP (#4906845001, Roche) was added to the lysis buffer if preservation of phosphorylated protein was required. Cell lysates were centrifuged at 21,000 RCF for 10 min, where the supernatant was collected and protein concentration was measured using the Micro BCA^TM^ Protein Assay Kit (#23235, Thermo Fisher Scientific). Total proteins were then added to Laemmli sample buffer (#161-0747, Bio-Rad) with 5% β-mercaptoethanol (#M3148, MilliporeSigma) and ran on an SDS-PAGE. The SDS-PAGE was transferred onto nitrocellulose membranes (#162-0115, Bio-Rad), which were blocked with 5% milk in 1x TBS-T (TBS with 0.1% Tween20 (#BP337-500, Fisher Scientific) for 1 hour at room temperature. The blots were then incubated with primary antibodies in 1% milk in 1x TBS-T overnight at 4°C. The following primary antibodies were used: anti-GABRA1 (mouse, 1:2000; #MAB339, MilliporeSigma), anti-Sodium Potassium ATPase (rabbit, 1:10000; #ab76020, Abcam), anti-ubiquitin (mouse, 1:2000; #14-6078-82, Invitrogen), anti-LC3b (rabbit, 1:2000; #NB100-2220, Novus Biologicals), anti-XBP1-s (rabbit, 1:1000; #12782S, Cell Signaling), anti-PERK (mouse, 1:1000; #5683, Cell Signaling), and β-actin (1:10000; #12004163, Bio-Rad). The next day, the blots were washed thrice with 1x TBS-T and incubated with secondary antibodies in 1% milk in 1x TBS-T for 1 hour at room temperature. The following secondary antibodies were used: goat anti-Rabbit IgG (1:10000; #A27036, Fisher Scientific) and goat anti-Mouse IgG (1:10000; #PIA28177, Fisher Scientific) or goat anti-Mouse AzureSpectra 700 (1:3000; #AC2129, Azure Biosystems). After incubation, the blots were washed twice with 1x TBS-T and twice more with 1x TBS. Lastly, SuperSignal^TM^ Pico PLUS or Femto chemiluminescent substrates (#PI34578, #PI34095, Fisher Scientific) were added to the blots to activate the HRP-conjugated antibodies. Blots were imaged using Azure Biosystems C600.

### Immunofluorescence

Cells were seeded on Matrigel (#54234, Corning)-coated coverslips and transfected according to cell culture protocol and fixed 48 hours after transfection with 4% paraformaldehyde (#15710, Electron Microscopy Sciences) in 1x DPBS for 15 minutes. Cells were then permeabilized by incubation in 0.2% saponin (#84510, MilliporeSigma) for 20 minutes at room temperature. However, when staining for surface proteins, no permeabilization was performed and saponin was not utilized at all. Then, non-specific binding sites were blocked using 10% BSA (#A-420-10, GoldBio) for 1 hour at room temperature. The cells were next incubated with primary antibodies in 0.2% saponin and 2% BSA at 4°C overnight. The following primary antibodies were used: anti-GABRA1 (mouse, 1:300; #MAB339, MilliporeSigma) and anti-Calnexin (goat, 1:300; #ab219644, Abcam). The next day, the cells were washed twice with 1x DPBS and incubated with secondary antibodies in 2% BSA for 1 hour at room temperature. The following secondary antibodies were used: Alexa 488 goat anti-mouse (1:500; #A11029, Invitrogen), Alexa-568 goat anti-rabbit (1:500; #A11036, Invitrogen), Alexa-568 donkey anti-mouse (1:500; #A11031, Invitrogen), Alexa-488 donkey anti-goat (1:500; #A11055, Invitrogen). After incubation, the cells were washed twice with 1x DPBS. Then, cells were incubated with DAPI dye (#HY-D0814, MedChemExpress) (1 μg/mL in 1x DPBS) for 10 minutes at room temperature. Coverslips were washed twice with 1x DPBS and mounted onto glass slides using Fluoromount-G^TM^ Mounting Medium (#00-4958-02, Thermo Fisher Scientific). Coverslips were imaged using an Olympus FV3000 microscope with a 60×1.40 numerical aperture oil objective.

### Automated patch clamp

Cells were seeded at 1.5 million cells per 10 cm plate and transiently transfected with α_1_, β_2_, and γ_2_ subunits (1:1:1) 24 hours after plating. Cells were lifted 48 hours post-transfection using Accutase (#A6964, Sigma Aldrich) centrifuged for 1 minute at 200 RCF and resuspended in Serum Free Media made of 293-SFM II (#11686029, Thermo Fisher), 25 mM HEPES (#15630-080, Gibco), 1% Penicillin-Streptomycin. Resuspended cells were left on a shaker for 30 minutes at room temperature. Then, cells were centrifuged at 200 RCF for 1 minute and resuspended in extracellular solution (ECS; 138 mM NaCl, 4 mM KCl, 1 mM MgCl2, 1.8 mM CaCl2, 5.6 mM Glucose, 10 mM HEPES). The intracellular solution (ICS; 70 mM KF, 60 mM KCl, 15 mM NaCl, 5 mM EGTA, 5 mM HEPES) alongside cells resuspended in ECS were applied to the Ionflux ensemble plate 16 (#910-0054, Fluxion Biosciences). Whole-cell GABA_A_R currents were then recorded using automated patch-clamping on the Ionflux Mercury 16 (Fluxion Biosciences), where the voltage was held at -60 mV and 100 μM GABA was applied for 3 seconds. Each ensemble recording enclosed the currents of 20 cells and was analyzed using Fluxion Data Analyzer.

### Endoglycosidase H (Endo H) assay

Cells were harvested and lysed according to our cell culture protocol. Then, 30 μg of cell lysate was incubated with Denaturing Buffer for 10 minutes at room temperature for the denaturation reaction. Then, the endoglycosidase H (endo H; #P0703L, NEBiolab) enzyme was incubated with the GlycoBuffer and denaturation solution for 1 hour at 37°C for the endo H digestion. For the negative control, no endo H was added, and for the positive control, GlycoBuffer, NP40, and PNGase F (#P0704L, NEBiolab) were added to the denaturation solution. After incubation, 4x Laemmli sample buffer with 5% β-mercaptoethanol was added to the solution and ran on an SDS-PAGE.

### Cycloheximide chase assay

Cycloheximide (100 μg/mL) was added to transfected HEK293T cells at multiple time points (0, 0.5, 1, 2, 3, 4 hours) to inhibit protein synthesis. Then, cells were lysed and ran on an SDS-PAGE. Bands were first normalized to β-actin and then normalized to the 0-hour time point.

### Surface Biotinylation

Cells were seeded on poly-*L*-lysine (#ICN15017710, Fisher Scientific) coated 6-cm dishes and transfected according to cell culture protocol. 48 hours after transfection, cells were washed with cold DPBS-CM (1x DPBS, 0.5 mM MgCl_2_, 1 mM CaCl_2_) and incubated with Sulfo-NHS-SS-Biotin (0.5 mg/mL; #A8005, Apex Bio) in DPBS-CM (1 mM CaCl2 and 0.5 mM MgCl2 dissolved in DPBS) for 30 minutes at 4°C. Cells were then incubated with 50 mM glycine in cold DPBS-CM for 5 minutes at 4°C, twice. Next, cells were washed twice with cold DPBS-CM and then incubated with 5 nM N-ethylmaleimide (NEM) in DPBS-CM for 15 minutes at room temperature. Lastly, the remaining solutions were aspirated from the plate and lysis buffer (2 mM DDM and 5 mM NEM in 1x TBS supplemented with complete protease inhibitor cocktail) was added for cell scraping. After scraping, the cell solution was then transferred to an Eppendorf tube and solubilized overnight at 4°C on an orbital rotor. The next day, lysates were centrifuged at 16,000 RCF for 10 minutes and the supernatant was then collected to be added to NeutrAvidin-conjugated agarose bead slurry (#29200, Thermo Fisher Scientific). The lysates were incubated with the slurry for 2 hours at 4°C on an orbital rotor. Samples were then centrifuged at 5,000 RCF for 30 seconds and the supernatant was collected as the cytosolic fraction. The beads were washed thrice with wash buffer (1% Triton X-100 in 1x TBS) and thrice more with wash buffer without detergent (1x TBS). Beads were pelleted by centrifugation at 5,000 RCF for 30 seconds and surface proteins were eluted by vortexing the beads for 20 minutes at room temperature in elution buffer (100 mM DTT and 6 mM urea in 2x Laemmli sample buffer). Beads were once more pelleted at 10,000 RCF for 60 seconds and the supernatant was collected to be ran on SDS-PAGE. The cytosolic fraction was added to Laemmli sample buffer and also ran on SDS-PAGE alongside the biotinylated fraction. Cytosolic bands were normalized to β-actin and surface bands were normalized to Na^+^/K^+^ ATPase.

### Quantitative reverse transcription polymerase chain reaction (qRT-PCR)

Cells were plated, transfected, and harvested according to our cell culture protocol. After lifting and centrifuging the cells, the pellet is resuspended in 1x DPBS, and total RNA was extracted from the cells using the High Pure RNA Isolation Kit (#11828665001, Roche). Then, cDNA was synthesized from 1 μg of total RNA using the iScript^TM^ cDNA synthesis kit (#1708891, Bio-Rad). PowerUp SYBR Green Master Mix (Applied Biosystems #A25776), primers (Integrated DNA Technologies; **Supplemental Table 1**) were mixed together for amplification (40 cycles of 15 s at 95 °C and 60 s at 60 °C) in QuantStudio 3 Real-Time PCR System (Applied Biosystems). Threshold cycle (C_T_) was determined from the PCR amplification plot, and the ΔC_T_ value was defined as: ΔC_T_□=□C_T_ (target gene) - C_T_ (housekeeping gene, *RPLP2*). The relative mRNA expression level of target genes of treated/variant cells was normalized to untreated/WT cells: Relative mRNA expression level□=□2 exp [- (ΔC_T_ (treated cells) - ΔC_T_ (untreated cells))]. Each data point involved technical triplicates.

## Statistical analysis

Quantification was performed using ImageJ, then plotted and analyzed using GraphPad Prism. Error bars were presented as mean ± standard error of the mean (SEM). Statistical significance was calculated using an unpaired t-test for two group comparisons, comparisons of more than two groups used an ANOVA followed by Dunnett’s test, and comparisons between groups on two independent variables used an unpaired t-test (**Supplemental Table 2**). Outliers were identified using a Grubbs’ test. A *p* value of less than 0.05 was considered statistically significant. *, *p* < 0.05; **, *p* < 0.01; ***, *p* < 0.001; ****, *p* < 0.0001.

## Supporting information

This article contains supporting information. Supporting information includes two supplemental tables and two supplemental figures.

## Acknowledgements

This work was supported by the National Institutes of Health (R01NS105789 and R01NS117176 to TM) and T32 training grant (GM008803 to MW).

## Author contributions

Conceptualization, MW, YW, and TM; Data curation: MW and YW; Formal analysis: MW, YW, and TM; Funding acquisition: MW and TM; Supervision: YW and TM; Writing – original draft: MW, YW, and TM; Writing – review & editing: MW, YW, JK, JP, and TM.

## Funding and additional information

The content is solely the responsibility of the authors and does not necessarily represent the official views of the National Institutes of Health.

## Supplemental Tables

**Table S1.**
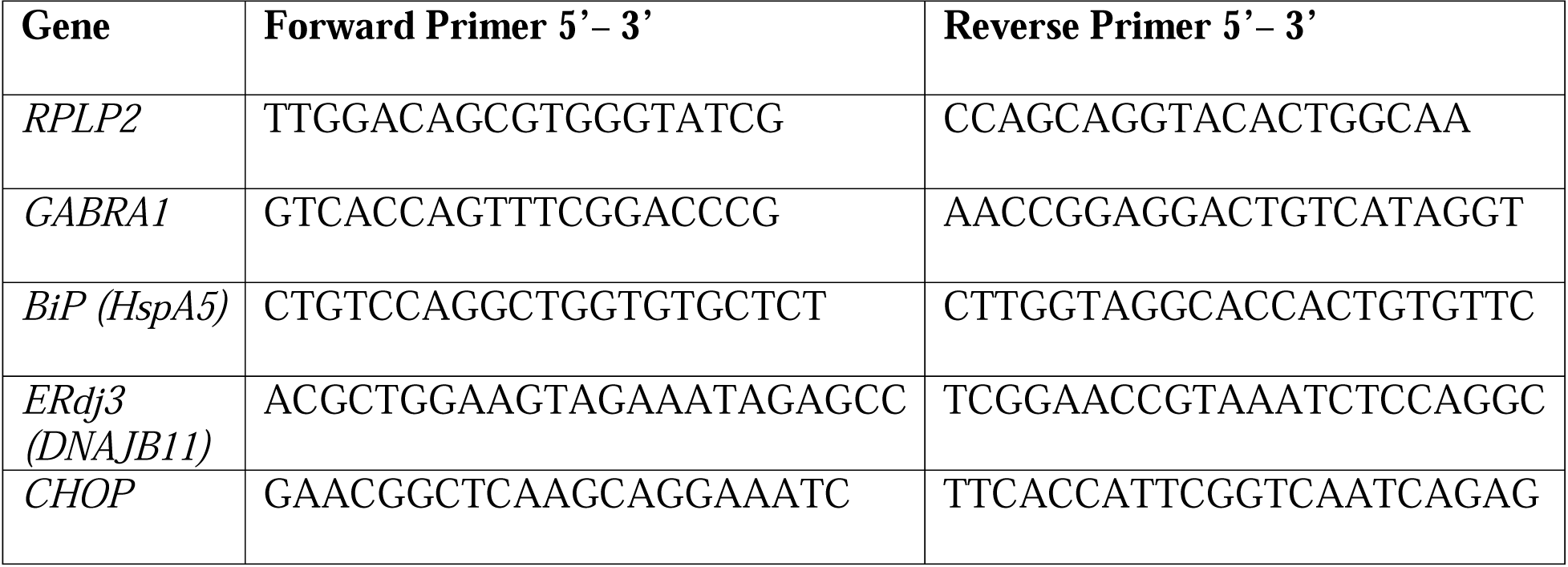
List of qPCR primers that were utilized in this study.

**Table S2.** List of statistical tests utilized in this study.

| Figure Panel | Assay | Statistical test | Adjusted P-value |
| --- | --- | --- | --- |
| Figure 1D | Total $\alpha_1$ expression | One-way ANOVA test, Dunnett's post-hoc test | WT vs K401<sub>fs</sub>; $p = 0.5060$<br>WT vs S326<sub>fs</sub>; $p = 0.0057$<br>WT vs V290<sub>fs</sub>; $p = <0.0001$<br>WT vs F272<sub>fs</sub>; $p = 0.1172$ |
| Figure 1E | Oligomer to monomer $\alpha_1$ ratio | One-way ANOVA test, Dunnett's post-hoc test | WT vs K401<sub>fs</sub>; $p = 0.1521$<br>WT vs S326<sub>fs</sub>; $p = 0.0045$<br>WT vs V290<sub>fs</sub>; $p = 0.0029$<br>WT vs F272<sub>fs</sub>; $p = 0.0018$ |
| Figure 2A | Surface biotinylation | One-way ANOVA test, Dunnett's post-hoc test | WT vs K401<sub>fs</sub>; $p = 0.0009$<br>WT vs S326<sub>fs</sub>; $p = 0.0018$<br>WT vs V290<sub>fs</sub>; $p = 0.0014$<br>WT vs F272<sub>fs</sub>; $p = 0.0004$ |
| Figure 2B | Immunocytochemistry –surface vs intracellular $\alpha_1$ | One-way ANOVA test, Dunnett's post-hoc test | WT vs K401<sub>fs</sub>; $p = <0.0001$<br>WT vs S326<sub>fs</sub>; $p = <0.0001$<br>WT vs V290<sub>fs</sub>; $p = <0.0001$<br>WT vs F272<sub>fs</sub>; $p = <0.0001$ |
| Figure 2C | Automated patch clamp recordings | One-way ANOVA test, Dunnett's post-hoc test | WT vs K401<sub>fs</sub>; $p = <0.0001$<br>WT vs S326<sub>fs</sub>; $p = <0.0001$<br>WT vs V290<sub>fs</sub>; $p = <0.0001$<br>WT vs F272<sub>fs</sub>; $p = <0.0001$ |
| Figure 3A | Endo H digestion | One-way ANOVA test, Dunnett's post hoc test | WT vs K401<sub>fs</sub>; $p = <0.0001$<br>WT vs S326<sub>fs</sub>; $p = <0.0001$<br>WT vs V290<sub>fs</sub>; $p = <0.0001$<br>WT vs F272<sub>fs</sub>; $p = <0.0001$ |
| Figure 3B | Immunocytochemistry –$\alpha_1$ and calnexin co- | One-way ANOVA test, | WT vs K401<sub>fs</sub>; $p = <0.0001$<br>WT vs S326<sub>fs</sub>; $p = <0.0001$ |
| | localization | Dunnett's post hoc test | WT vs V290<sub>fs</sub>; $p = 0.0041$<br>WT vs F272<sub>fs</sub>; $p = 0.0405$ |
| Figure 4A | Proteasomal vs lysosomal degradation | One-way ANOVA test, Dunnett's post-hoc test | WT: DMSO vs BafA1; $p = 0.0038$<br>S326<sub>fs</sub>: DMSO vs BafA1; $p = 0.0158$<br>S326<sub>fs</sub>: DMSO vs MG132; $p = 0.0184$<br>F272<sub>fs</sub>: DMSO vs BafA1; $p = 0.0020$ |
| Figure 4B | qPCR analysis for NMD | Unpaired t-test | S326<sub>fs</sub>; $p = 0.0166$<br>V290<sub>fs</sub>; $p = <0.0001$<br>F272<sub>fs</sub>; $p = 0.0090$ |
| Figure 5A | qPCR analysis for UPR activation | One-way ANOVA test, Dunnett's post-hoc test | ERdj3:<br>WT vs K401<sub>fs</sub>; $p = 0.3256$<br>WT vs S326<sub>fs</sub>; $p = 0.2819$<br>WT vs V290<sub>fs</sub>; $p = 0.0009$<br>WT vs F272<sub>fs</sub>; $p = 0.0211$<br>BiP:<br>WT vs K401<sub>fs</sub>; $p = 0.0693$<br>WT vs S326<sub>fs</sub>; $p = 0.1084$<br>WT vs V290<sub>fs</sub>; $p = <0.0001$<br>WT vs F272<sub>fs</sub>; $p = 0.0294$<br>CHOP:<br>WT vs K401<sub>fs</sub>; $p = 0.0207$<br>WT vs S326<sub>fs</sub>; $p = 0.1448$<br>WT vs V290<sub>fs</sub>; $p = 0.0011$<br>WT vs F272<sub>fs</sub>; $p = 0.2414$ |
| Figure 5B | IRE1 inhibition | Unpaired t-test | WT: DMSO vs KIRA6; $p = 0.097955$<br>K401<sub>fs</sub>: DMSO vs KIRA6; $p = 0.00026$<br>S326<sub>fs</sub>: DMSO vs KIRA6; $p = 0.000176$<br>V290<sub>fs</sub>: DMSO vs KIRA6; $p = 0.045941$<br>F272<sub>fs</sub>: DMSO vs KIRA6; $p = 0.000462$ |
| Figure 5C | PERK inhibition | Unpaired t-test | WT: DMSO vs GSK; $p = 0.080790$<br>K401<sub>fs</sub>: DMSO vs GSK; $p = 0.001282$<br>S326<sub>fs</sub>: DMSO vs GSK; $p = 0.023712$<br>V290<sub>fs</sub>: DMSO vs GSK; $p = 0.168432$<br>F272<sub>fs</sub>: DMSO vs GSK; $p = 0.264301$ |

## Supplemental figure legends

**Supplemental Figure S1.**
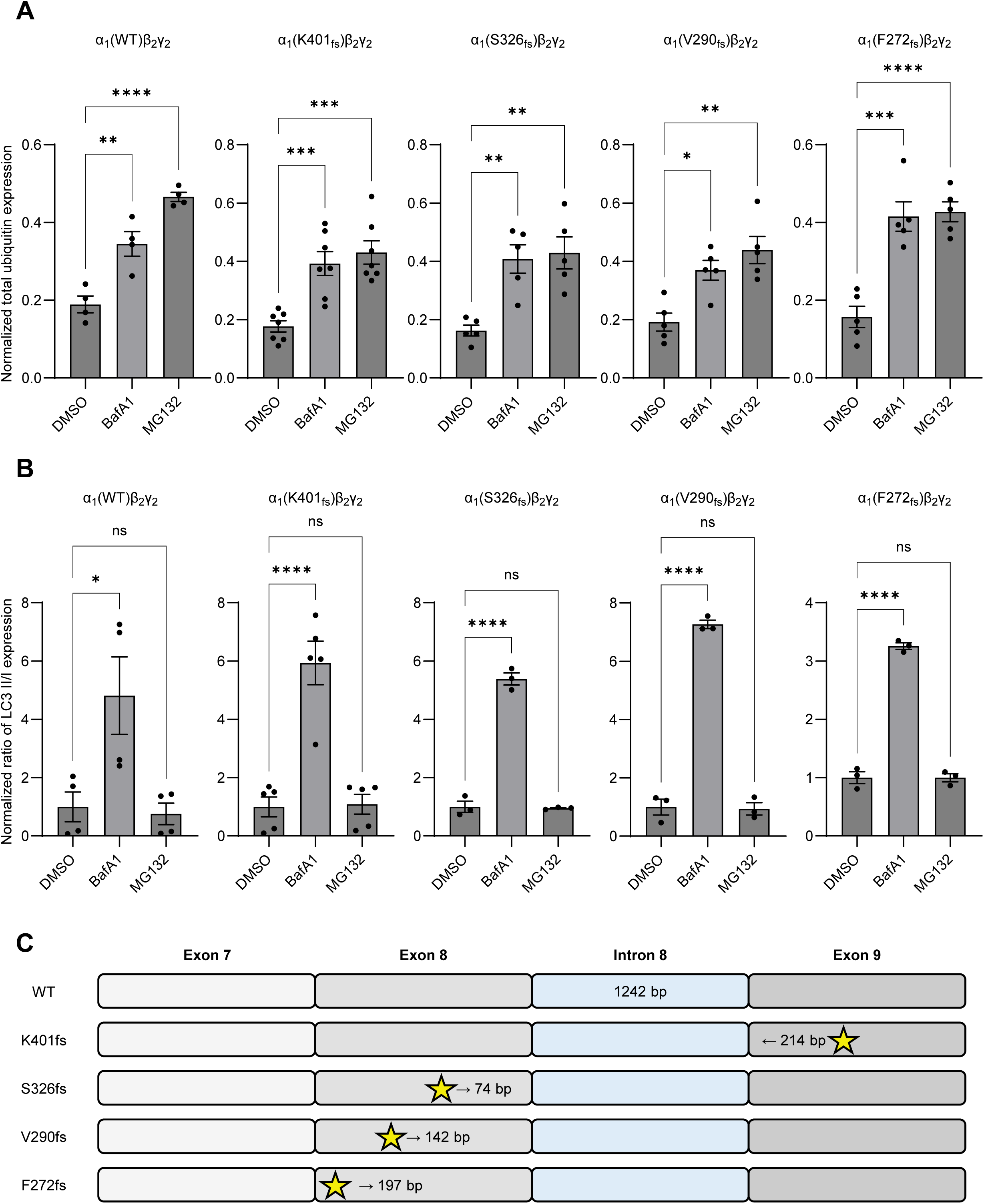
Inhibition of the proteasome or the lysosome. To inhibit proteasomal and lysosomal degradation, MG132 (200 nM, 24 hrs) and BafA1 (20 nM, 24 hrs) were applied, respectively. Total α_1_ proteins were extracted 48 hours after transfection, followed by SDS-PAGE and Western blot to detect the ubiquitin protein (**A**) and the LC3b isoforms (**B**). β-actin was used as a loading control. Data is presented as mean ± SEM. (**C**) Cartoon of minigene constructs of human *GABRA1* containing intron 8, showing the stop codons resulting from the K401, S326, V290, and F272 deletions (indicated by the star). One-way ANOVA followed by Dunnett’s test was used for statistical analysis. *, *p* < 0.05; **, *p* < 0.01; ***, *p* < 0.001, ****, *p* < 0.0001.

**Supplemental Figure S2.**
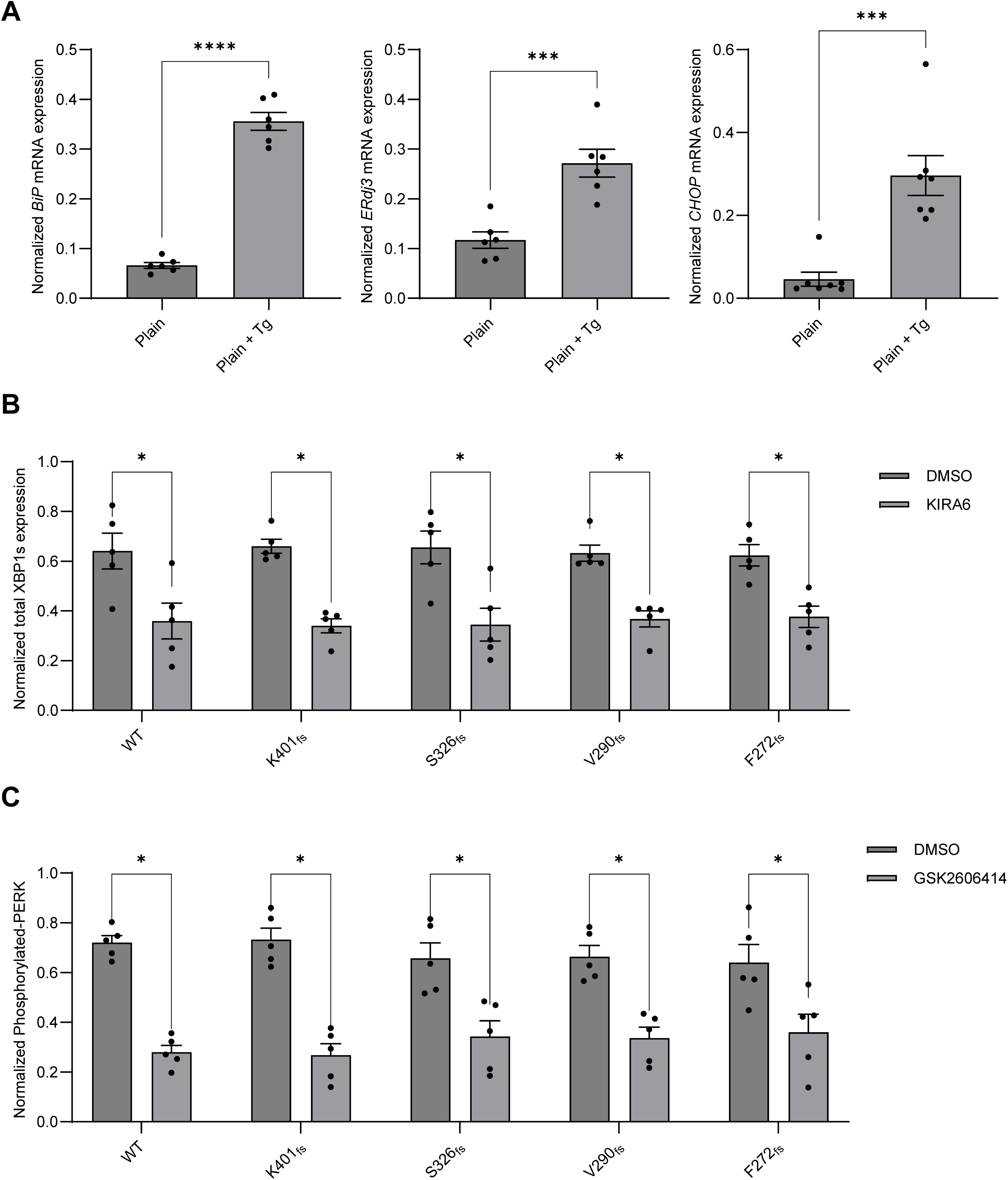
Effect of α_1_ variants on the UPR. (**A**) Thapsigargin (Tg, 1 µM, 6 hrs) was applied to HEK293T cells as a positive control of UPR activation. Relative mRNA expression of UPR targets, including BiP, ERdj3, and CHOP for negative (plain) and positive control (plain + Tg) samples (n=5-6). (**B**) KIRA6 (1 µM, 24 hrs), an IRE1 inhibitor, was applied to HEK293T cells expressing WT or frameshift α1 mutants. Total proteins were extracted 48 hours after transfection, followed by SDS-PAGE and Western blot to detect XBP1-s and confirm IRE1 inhibition (n=5). (**C**) GSK2606414 (0.5 µM, 24 hrs), a PERK inhibitor, was applied to HEK293T cells expressing WT or frameshift α1 mutants. Total proteins were extracted 48 hours post transfection, followed by SDS-PAGE and Western blot to detect the PERK and phosphorylated PERK (n=5). Proteins bands were quantified using ImageJ. β-actin was used as a loading control. Data is presented as mean ± SEM. Unpaired T-test was used for two group comparisons. *, *p* < 0.05; ***, *p* < 0.001; ****, *p* < 0.0001.

